# A Collection of 2,280 Public Domain (CC0) Curated Human Genotypes

**DOI:** 10.1101/127241

**Authors:** Richard J. Shaw, Manuel Corpas

**Affiliations:** Repositive Ltd, Future Business Centre, Cambridge, UK

## Abstract

Cheap sequencing has driven the proliferation of big human genome data aggregation consortiums, providing extensive reference datasets for genome research. These datasets, however, may come with restrictive terms of use, conditioned by the consent frameworks within which individuals donate their data. Having an aggregated genome dataset with unrestricted use, analogous to public domain licensing, is therefore unusually rare. Yet public domain data is tremendously useful because it allows freedom to perform research with it. This comes with the price of donors surrendering their privacy and accepting the associated risks derived from publishing personal data. Using the Repositive platform (https://repositive.io/?23andMe), an indexing service for human genome datasets, we aggregated all deposited files in public data sources under a CC0 license from 23andMe, a leading Direct-to-Consumer genetic testing service. After downloading 3,137 genotypes, we filtered out those that were incomplete, corrupt or duplicated, ending up with a dataset of 2,280 curated files, each one corresponding to a unique individual. Although the size of this dataset is modest compared to current major genome data aggregation projects, its full access and licensing terms, which allows free reuse without attribution, make it a useful reference pool for validation purposes and control experiments.

## Background & Summary

The availability of personal genome Direct-to-Consumer (DTC) tests has been fuelled by companies like 23andMe, which makes it easy for users to access their personal genotype data for an affordable price. Despite recent improvements in cost and availability of Next Generation Sequencing (NGS) technology for DTC use, array genotyping is still a viable option [1], particularly when users want to have a whole genome screen of known Single Nucleotide Variations (SNVs) for a cheap price. For about $100 one can buy a DTC test and have a satisfying state-of-the-art analysis of ancestry and genetic health risk [2]. 23andMe has been able to capitalise on the curiosity of many customers from around the world, generating one of the most extensive private genome data ecosystems in the world [3]. The genotype data aggregated from 23andMe customers, combined with phenotype information from questionnaires, has already been proven to be an effective way to discover new markers, e.g. [4–6].

Using the genotype data from 23andMe as a gateway for personal/recreational genomics also has other advantages. The format of the different SNP array versions (although the number of tested SNVs may vary) remains constant and relatively easy to handle: the size of the genotype files is in the order of tens of MB, which is more manageable than the GB sizes of Next Generation Sequencing (NGS) files. 23andMe genotype files are also a useful proxy for understanding the individual’s main genetic features.

In this study we use the Repositive platform [7] (https://repositive.io/?23andMe), which indexes human genomic datasets from all major and known genome data repositories, to gather the greatest possible open access 23andMe set of individual genotypes. We took only 23andMe genotypes that have been deposited in public archives and have no restriction of use, i.e. a CC0 license (Table 1). These archives include (in order of greatest contribution): openSNP [8], the Personal Genome Project [9], Open Humans [10], Genomes Unzipped [11], the Corpasome [12] and Stephen Keating’s data source [13]. Any genotype from these data sources can be copied, modified, distributed and used, even for commercial purposes, all without having to ask permission.

**Table 1:** Summary of licenses and their provenance for all data used for this study. All of them are in the public domain (CC0) [14]. This means that the data referenced in this study can be used, distributed or modified, even for commercial purposes, without needing to ask permission to the individuals from whom this data originated.

| 23andMe Data Source | License | Terms of Use |
| --- | --- | --- |
| openSNP | CC0 | https://www.iubenda.com/privacy-policy/641811 |
| PGP | CC0 | http://www.personalgenomes.org/tos |
| Open Humans | CC0 | https://www.openhumans.org/community-guidelines/#public-data |
| Genomes Unzipped | CC0 | http://genomesunzipped.org/data |
| Corpasome GRCh37 | CC0 | https://figshare.com/articles/23andMe_hg37/4491215 |
| Stephen Keating | CC0 | http://stevenkeating.info/main.html |

We downloaded every open access public domain 23andMe genotype available in every public repository currently indexed by Repositive, curating and bundling them into a single dataset. This amounted to 3,137 genotype files that, after curation, were reduced to 2,280 genotypes.

These 2,280 genotypes constitute an open access dataset of high utility in the personal genomics field. Although a number of resources are available that provide genome-based SNV data at a population level (e.g., 1000K genomes [15], ExAC [16], GnomAD [17], UK10K [18], PGP [9]), to date there is no a single repository that meets all the stated characteristics our proposed aggregation dataset provides: full genotype data, open access, unrestricted use, derived from the same source at the scale of thousands. Hence we believe this dataset will prove a valuable resource for the genome research community, both academic and industry, who will be able to benefit from a curated cohort of genotypes.

## Methods

In order to aggregate all open access 23andMe genotypes in the public domain we used the Repositive platform [7] (https://repositive.io/?23andMe). Repositive is an indexing service that catalogues all known human genome datasets in public sources and repositories throughout the internet. Repositive stores both the metadata which describes the deposited dataset, and the link through which the dataset is accessed. A total of 3,137 links from the Repositive platform matched a potential 23andMe entry (complete list of links available in Suppl. File 1^1^). These links pointed to 6 sources: 2,318 from openSNP, 514 from the Personal Genome Project (PGP), 286 from Open Humans, 13 from Genomes Unzipped, 5 from the Corpasome [19] and 1 from Stephen Keating’s source [20]. We then downloaded all the files and proceeded with their curation:

1. Among the 3,137 downloaded files we found 37 that were non-text files: e.g., PDFs or image files.
2. For the remaining 3,100 files we discarded 210 files (168 VCFs and 42 of unknown format)
3. This left us a total of 2,890 files:

a. 2,484 for build GRCh37
b. 378 for build 36
c. 28 for unknown build
4. From the 2,484 that mapped to the GRCh37 build, we discarded 87 that had less than 500k SNP rows, giving us a total of 2,397 23andMe text files. The Corpasome 23andMe files (corresponding to 5 individuals from the Corpas family) were mapped to build 36. Since we had direct access to their 23andMe account, we downloaded them directly from 23andMe, this time mapped to build 37. This produced a total of 2,402 GRCh37 23andMe text files.
5. We further refined the 2,402 GRCh37 text files, extracting the consensus SNP positions from all autosomes (chromosome 1-22). This yielded a total of 445,734 SNPs. Using the R packages gdsfmt and SNPRelate[21], the 445,734 SNPs were further reduced to 63,486 by linkage pruning.
6. We then created a Principal Component Analysis (PCA) to cluster the 23andMe data from collected individuals into populations. The resulting PCA showed 141 very clear duplications.
7. Note also that the 141 duplications do not necessarily imply 2402 - 141 unique samples. Since some samples were multiply duplicated, there were actually 2280 unique genotypes (the Corpasome is included).

### Code availability

A python script that takes the 23andMe URLs and downloads them is available: https://drive.google.com/file/d/0B8yXU9SkT3ftZHk4ejRCUUtlVm8/view

## Data Records

For version 1 and 2 of their SNP chip 23andMe used a customised Illumina Hap550+ array. Version 3 is based on a customised Illumina OmniExpress+ array and version 4 uses a custom Illumina HumanOmniExpress-24 format chip. The downloaded raw data file is a zipped text file 5 MB to 30 MB in size in bed format [22], with a header describing the date in which the file was downloaded and the human reference assembly build used as coordinates for the mapping of the SNVs tested. The file itself provides the SNPdb ID for each SNV [23], the chromosome and position in the mapped assembly and the genotype, including both alleles. Figure 2 shows a snippet of the 23andMe genotype bed file from one of us downloaded on “Tue Dec 13 09:12:17 2016”.

**Figure 1:**
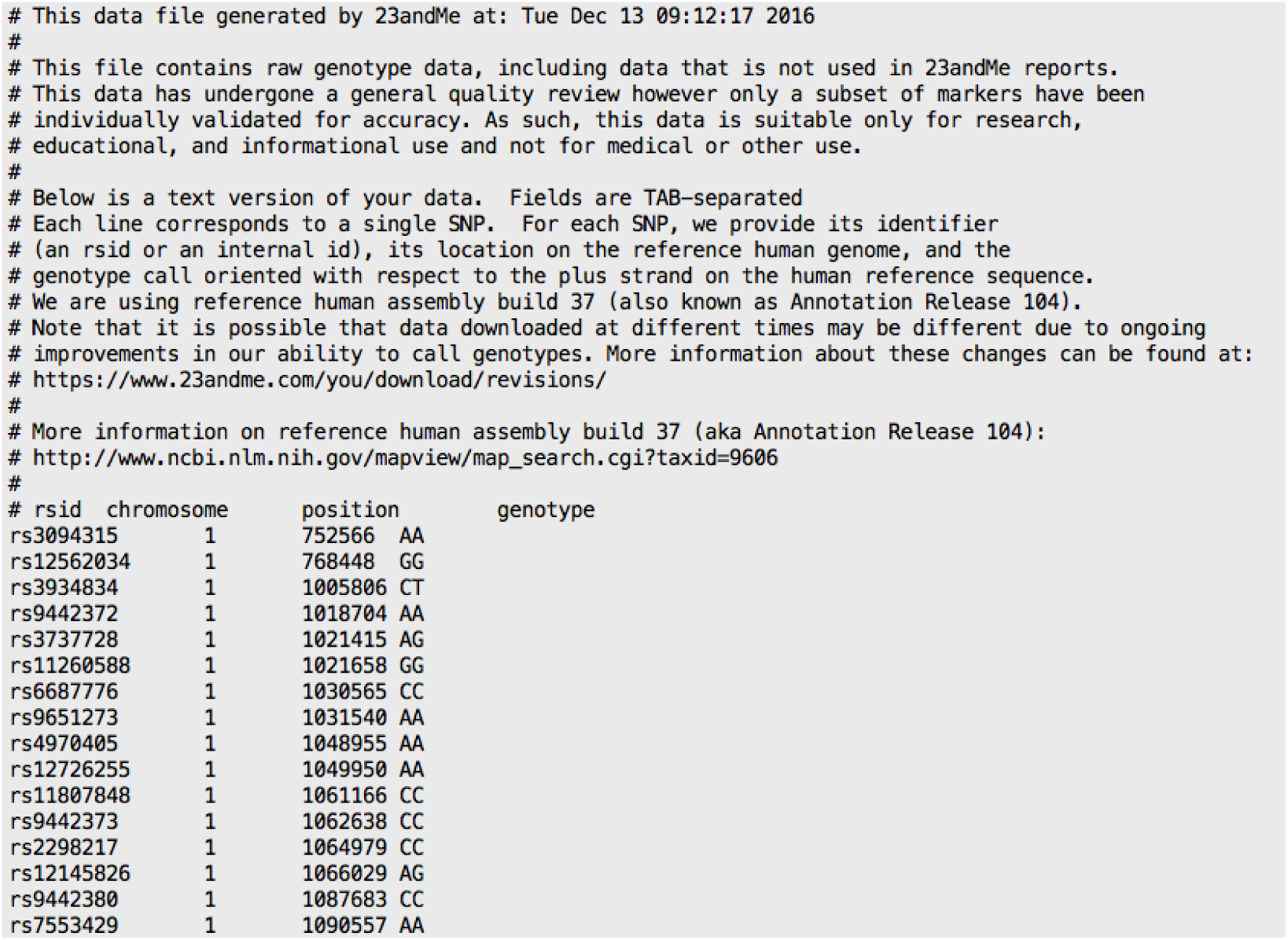
A screenshot of the top of a 23andMe raw data genotype showing the date on which it was downloaded, the assembly (human assembly build 37) and, for each SNV analysed, its SNV ID (RSID), its location (chromosome and position) and the genotype for both alleles at that position.

**Figure 2:**
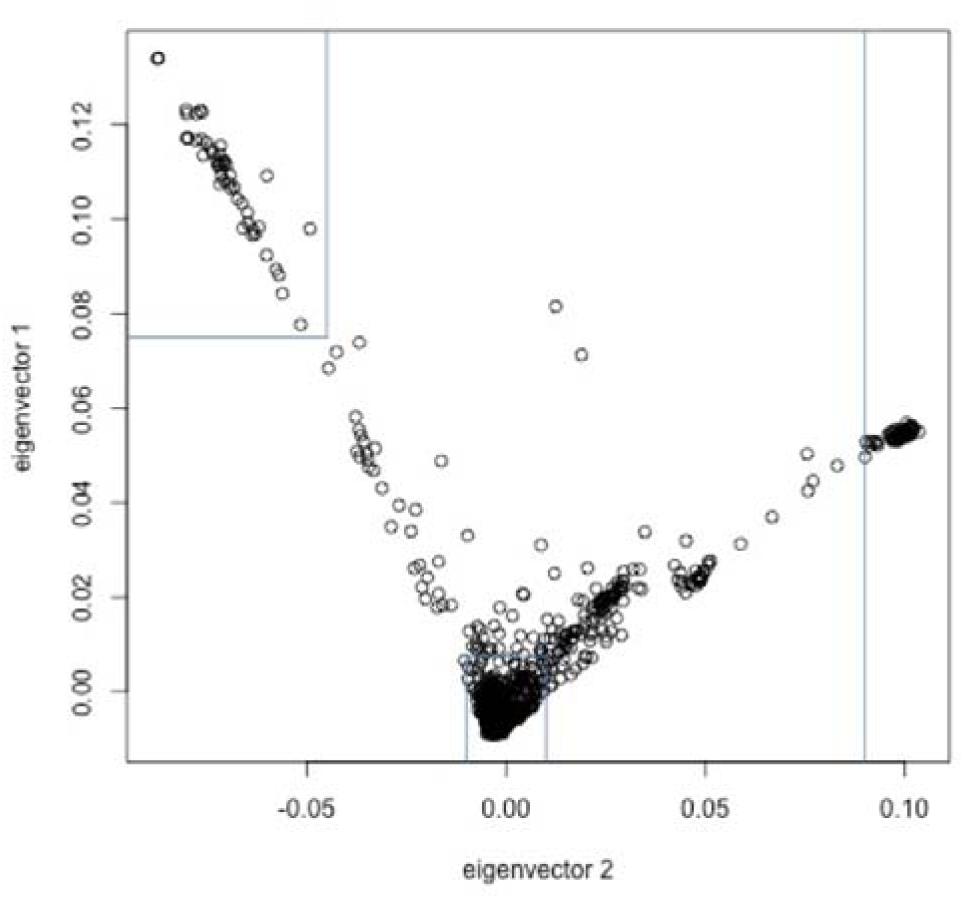
Principal Component Analysis (PCA) of 2,402 23andMe data files from individuals who have openly shared their genotypes in public repositories. We found, by looking at their PCA values, that there are 141 that are duplicated.

Table 2 summarises the number of links for each data source, the number of text files downloaded and the number of 23andMe files mapped to the build 37 with more than 500k SNPs each.

**Table 2:** Summary of the total number of links obtained through the Repositive platform and their sources. The data source from which we get the greatest number of links for 23andMe data files is openSNP. A process of curation was carried out to select only 23andMe files that map to build GRCh37 and have >500k SNPs.

| Source | Total # Links | Downloaded Text Files | GRCh37 23andMe Text Files with >500k SNP |
| --- | --- | --- | --- |
| openSNP | 2,318 | 2,296 | 1,947 (81%) |
| PGP | 514 | 499 | 316 (13%) |
| Open Humans | 286 | 286 | 133 (5.5%) |
| Genomes Unzipped | 13 | 13 | 0 |
| Corpasome | 5 | 5 | 5 (0.2%) |
| Stephen Keating | 1 | 1 | 1 (0.04%) |
| Total | 3,137 | 3,100 | 2,402 |

## Data Availability

The 2,280 curated set of links is available:

https://drive.google.com/file/d/0B8yXU9SkT3ftR3pIbW81cDhrc2s/view

The data can be downloaded using the python script provided at the link supplied earlier.

### Technical Validation

One common problem when aggregating genome data from different data sources is duplication. In this case, users wishing to share their genotype data may have submitted their 23andMe file to more than one open access repository.

We created a Principal Component Analysis (PCA) to cluster the 23andMe data from the 2,402 collected 23andMe individuals with more than 500k SNPs mapped to GRCh37 (Table 2). The resulting PCA clusters are shown in Figure 2. From the PCA, we were able to identify 141 very clear duplications. There were some further cases of very similar (but not identical) values but there were no obvious cases of siblings or other relatives from the metadata associated to these 23andMe files.

## Footnotes

1 https://docs.google.com/document/d/1R7zoqw7p1xwLxOpTTv0vWf88gZatiVke2k8M0J0sr8/edit

## Acknowledgements

We are grateful to the public repositories that make it possible for users to upload their genomic datasets at no cost.

## Author contributions

MC conceived the study and wrote the paper.

RS performed the curation and validation of the data.

